# Thousands of large-scale RNA sequencing experiments yield a comprehensive new human gene list and reveal extensive transcriptional noise

**DOI:** 10.1101/332825

**Authors:** Mihaela Pertea, Alaina Shumate, Geo Pertea, Ales Varabyou, Yu-Chi Chang, Anil K. Madugundu, Akhilesh Pandey, Steven L. Salzberg

## Abstract

We assembled the sequences from 9,795 RNA sequencing experiments, collected from 31 human tissues and hundreds of subjects as part of the GTEx project, to create a new, comprehensive catalog of human genes and transcripts. The new human gene database contains 43,162 genes, of which 21,306 are protein-coding and 21,856 are noncoding, and a total of 323,824 transcripts, for an average of 7.5 transcripts per gene. Our expanded gene list includes 4,998 novel genes (1,178 coding and 3,819 noncoding) and 97,511 novel splice variants of protein-coding genes as compared to the most recent human gene catalogs. We detected over 30 million additional transcripts at more than 650,000 sites, nearly all of which are likely to be nonfunctional, revealing a heretofore unappreciated amount of transcriptional noise in human cells.

## Introduction

Scientists have been attempting to estimate the number of human genes for more than 50 years, dating back to 1964 ^1^. In the decade preceding the initial publication of the human genome, multiple estimates were made based on sequencing of short messenger RNA fragments, and most of these estimates fell in the range of 50,000–100,000 genes ^2-5^. When the human genome was published in 2001, the estimates of the gene count were dramatically lower, with one paper reporting 30,000-40,000 genes ^6^ and the other 26,588 plus ∼12,000 genes with “weak supporting evidence” ^7^. As the genome was gradually made more complete and the annotation improved, the number continued to fall; when the first major genome update was published in 2004, the estimated gene count was revised to 20,000–25,000 ^8^. Later efforts suggested that the true number of protein-coding genes was even smaller: a 2007 comparative genomics analysis suggested 20,500 ^9^, and a proteomics-based study in 2014 estimated 19,000 ^10^.

One striking feature of most attempts to catalog all human genes is their lack of precision. Most estimates have only 1-2 significant digits, indicating major uncertainty about the exact number. As we reported in 2010, the estimates of the human gene count at that time averaged ∼22,500 genes ^11^. As of late 2017, one of the most reliable catalogs of human genes, the curated reference set from NCBI’s RefSeq database ^12^, contained 20,054 distinct protein-coding genes, and another widely used human gene catalog, Gencode ^13^, contained 19,817. The international CCDS database, an ongoing effort to identify all human and mouse genes ^14^, listed 18,894 human protein-coding genes in March 2018 (release 20).

The human gene list has a tremendous impact on biomedical research. A huge and still-growing number of genetic studies depend on this list, for example:

- Exome sequencing projects use exon capture kits that target most “known” exons. Any exons that are not listed in standard human annotation are ignored.
- Genome-wide association studies (GWAS) attempt to link genetic variants to nearby genes, relying on standard catalogs of human genes.
- Many software packages that analyze RNA sequencing (RNA-seq) experiments, which measure gene expression, rely on a database of known genes and cannot measure genes or splice variants unless they are included in the database.
- Efforts to identify cancer-causing mutations usually focus on mutations that involve known genes, ignoring mutations that occur in other regions.

These and other examples encompass thousands of experiments and an enormous investment of time and effort. The creation of a more complete, accurate human gene catalog will have an impact on many of these studies. For example, exome sequencing studies targeting Mendelian diseases, which should be the easiest diseases to solve, have reported diagnostic success in only about 25% of cases ^15,16^, perhaps because many exons and genes are excluded from exome capture kits. A better gene list may also help to explain the genetic causes of the many complex diseases that have thus far remained largely unexplained, despite hundreds of large GWAS and other experiments.

As part of the creation of a human gene list, we must first define what is meant by the term “gene.” During the Human Genome Project, most efforts to estimate and annotate genes focused on protein-coding genes; i.e., regions of the genome that are transcribed into RNA and then translated into proteins. At the time, most scientists assumed that non-coding genes represented only a very small portion of the functional elements of the human genome, and that most RNA genes (e.g., transfer RNAs and ribosomal RNA genes) were already known. A few years after the initial publication of the human genome, though, scientists began to uncover a large and previously-unappreciated complement of long noncoding RNA genes, lncRNAs ^17,18^, which quickly grew to include thousands of novel genes. These genes have a wide range of functions that are just as vital to human biology as many protein-coding genes ^19^, and any comprehensive list of human genes should include them.

Thus for the purposes of our study, genes will include any interval along the chromosomal DNA that is transcribed and then translated into a functional protein, *or* that is transcribed into a functional RNA molecule. By “functional” we mean to include any gene that appears to perform a biological function, even one that might not be essential. We recognize that the proper determination of function can be a lengthy, complex process, and that at present the function of many human genes is unknown or only partially understood. Our definition intentionally excludes pseudogenes, which are gene-like sequences that may arise through DNA duplication events or through reverse transcription of processed mRNA transcripts. Following previous conventions ^11^, when multiple proteins or RNA genes are produced from the same region through alternative splicing or alternative transcription initiation, we will count these variants as part of a single gene. Our total gene count, therefore, corresponds to the total number of distinct chromosomal intervals, or loci, that encode either proteins or noncoding RNAs; in addition we report the total number of gene variants, which includes all alternative transcripts expressed at each locus.

## Results

The basis for our human gene catalog is a new analysis of a large, comprehensive survey of gene expression in human tissues, the genotype-tissue expression (GTEx) study, which included samples from dozens of tissues collected from hundreds of individuals ^20^. All of these samples were subjected to deep RNA-sequencing, with tens of millions of sequences (“reads”) captured from each sample, in an effort to measure gene expression levels across a broad range of human cell types. This exceptionally large set of transcript data–just under 900 billion reads–provided an opportunity to construct a new, comprehensive set of human genes and transcripts. We accomplished this by assembling all of the samples, merging the results, and applying a series of computational filters to remove transcripts with insufficient evidence.

During the Human Genome Project, the gold standard for identifying a gene was evidence that it was transcribed into messenger RNA. This was the basis for the first large-scale effort to capture and catalog human genes ^21^ and for many subsequent efforts. However, over time it has become clear that the mere fact that a region of the genome is transcribed is insufficient to prove that it has a function, especially in light of evidence that random mutations can easily create transcriptional start sites ^22^. A second, arguably more powerful piece of evidence that a sequence is a gene is evolutionary conservation: if a protein sequence has been conserved in other species, this provides strong evidence that the sequence provides a useful function; i.e., that it is a gene. A third line of evidence is reproducibility: if we observe a transcript in multiple samples from multiple individuals, then it is less likely be the result of random transcription. We used each of these lines of evidence in constructing the new gene catalog.

## Novel genes and transcripts

We assembled all 9,795 RNA-seq samples from the GTEx collection (see Methods) and removed all transcripts that overlapped with known protein-coding genes, noncoding genes, or pseudogenes from RefSeq ^12^ or Gencode ^13^. This process generated 5,081,171 novel transcripts at 668,018 loci, where “novel” means that the transcripts did not overlap any annotated genes in the RefSeq or Gencode databases. We then used a variety of criteria, described below, to eliminate transcripts due to “noise” ^23^; i.e., transcripts produced by low-level transcriptional activity that appears to have no functional utility.

This noise is so ubiquitous that some computational methods for analyzing RNA-seq experiments automatically impose a threshold below which they will not report a transcript, even if reads are present ^24,25^. We also eliminated novel transcripts with no introns, which we assumed to be either noise or pseudogenes, unless they had high-expression levels and contained a potential protein-coding gene, as detailed below. Out of the 5,081,171 novel transcripts, only 139,289 (2.7%) in 41,979 (6.3%) distinct loci had at least one intron.

## Protein-coding genes

To identify novel protein-coding genes, we eliminated transcripts based on a series of relatively strict criteria designed to remove noise, pseudogenes, and alignment artifacts. For each transcript, we used blastx ^26^ to search all open reading frames against all mammalian proteins in GenBank and in UniProtKB/Swiss-Prot to determine whether any were conserved in other species or elsewhere in the human genome. We imposed the following criteria before determining that any transcript encodes a protein:

- The transcript must contain at least one intron and it must have expression level TPM>1 in at least one tissue, or alternatively it may be a single-exon transcript with expression level at least as high as the outliers for known transcripts, defined as TPM>13.87 (see Supplementary Materials).
- The transcript must not be contained in another transcript, unless it is expressed in more samples than all transcripts that contain it.
- The length of the open reading frame (ORF) must be at least 60 amino acids.
- The ORF cannot overlap known LINE or LTR repeat elements, or overlap ribosomal RNA genes.
- The BLAST e-value of the best protein alignment must be 10^−15^ or smaller.
- If the predicted protein matches another protein, the length of the ORF must be at least 75% of the length of the matching protein (in order to eliminate pseudogenes, which tend to be truncated).
- If predicted transcripts are in conflicting loci (i.e. overlapping transcripts on opposite strands) we only keep those that align to proteins with known functions.

After applying these filters, we were left with 1,178 novel protein-coding genes that included 1,335 protein-coding transcripts (Supplementary Files S1 and S2). 601 of the novel protein-coding genes (654 transcripts) had their best match to non-human proteins in either GenBank or UniProtKB/Swiss-Prot. All but 3 genes had alignments to mammalian proteins in GenBank, while 580 also had protein hits to UniProtKB/Swiss-Prot. (The UniProtKB/Swiss-Prot database is manually annotated and thus higher quality but less complete than GenBank ^27^.) Combining the 1,178 new genes with the 20,054 from RefSeq yielded a total of 21,232 protein-coding genes.

We also considered using stricter criteria: if we strengthen our first criterion in the list above and require that a novel multi-exon protein occurs in at least 10 samples (rather than one sample) with an average TPM>1, the number of novel protein coding genes remained after our initial filters above would be reduced from 1,178 to 469. We retained the 709 novel genes that failed this stricter test, but labeled them as low-confidence in the CHESS database.

Figure 1 illustrates one of the novel genes, CHS.7402, discovered by this process. This four-exon gene occurs on chromosome 10 and spans the range 122,657,410–122,679,509, approximately 14 Kb downstream from the nearest known gene, DMBT1. It is highly conserved in multiple other species, with the closest homologs in macaques (94% identical over the full length of the protein, BLAST e-value 1e-38), followed by marmoset, capuchin, ass, Przewalski’s horse, rhinoceros, wild boar, and others (**Figure 1**).

**Figure 1:**
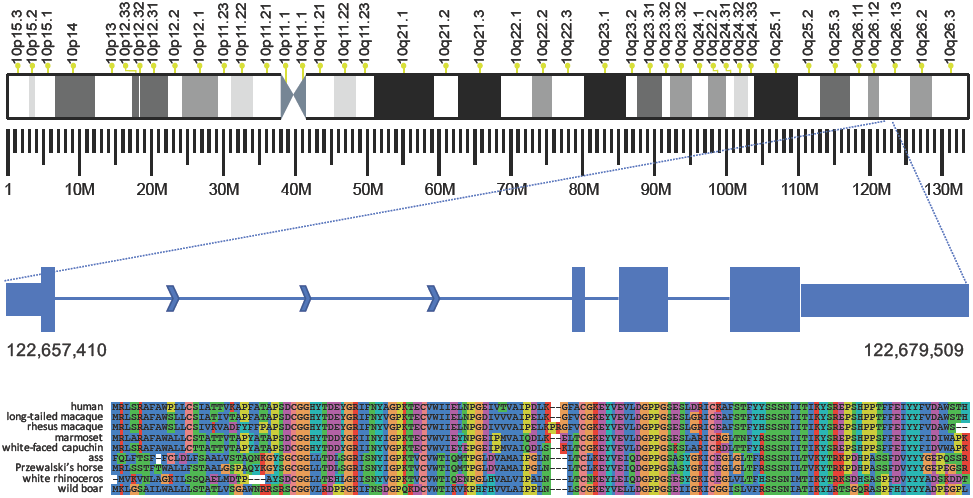
One of 1,587 new protein-coding genes (CHS.7402) discovered in this study. This 4-exon gene occurs on the forward strand of chromosome 10 at the coordinates shown. The exon lengths are 134, 30, 136, and 663 bp (left to right), with the narrower rectangles indicating the 5’ and 3’ UTR regions. The intron lengths (not shown to scale) are 18098, 1086, and 1956 bp. The sequence alignment at the bottom shows, top to bottom, the protein sequences from CHS.7402, long-tailed macaque, rhesus macaque, marmoset, white-faced capuchin, ass, Przewalski’s horse, white rhinoceros, and wild boar. The full-length human protein sequence is shown.

Interestingly, we found no homologous proteins annotated in primates much more closely related to humans such as chimpanzee, gorilla, and orangutan. We searched the transcript sequence of CHS.7402 against the DNA of chimpanzee (*Pan troglodytes*) and found that the sequence matches nearly perfectly, at 97% identity over its entire length, and that chimpanzee also has four exons. Thus the gene is clearly present, though un-annotated, in *Pan troglodytes*. This illustrates a broader problem with gene annotation: when annotation is created for a new genome, which is typically done through a highly automated process, previously annotated genes from other species provide critical evidence to support the new annotation. Thus if a gene is missing from the human annotation, it may be omitted from the annotation of other species, especially close human relatives. Multiple sequence alignments for additional novel CHESS proteins are shown in Suppl. Figures S6-S11.

We then evaluated the 15,779 lncRNA genes in RefSeq to determine if any of these might instead be protein-coding genes. From all RefSeq lncRNAs, we selected those containing an ORF at least 180 bp (60 amino acids) long and searched these against the mammalian protein database. After excluding read-through transcripts, we found 2,762 potential protein sequences that matched a mammalian protein with an E-value of 10^−15^ or less, and were at least 75% as long as the best matching protein, using the same criteria as used above for novel protein-coding genes.

Because this step was intended only to recover mis-annotated lncRNAs that are likely to be proteins, we retained only those genes for which the best-matching protein had a named function; i.e., we excluded any lncRNA whose best hit was to a protein annotated as hypothetical, unknown, or uncharacterized (see Methods). After removing hits to proteins that had no associated function, we were able to rescue 53 genes containing 85 transcripts that passed all our criteria for protein-coding regions. Adding these 53 new protein-coding genes to our total, the number of protein-coding genes in the human gene catalog increased to 21,285.

Finally, we considered genes from the curated Gencode database ^13^ (releases 25 and 27) that were annotated as “known” protein-coding genes but were missing from RefSeq. Based on proteins that remained in release 27 of Gencode (see Supplementary File S3), we found 26 more protein-coding genes, for a total of 21,311. Finally we substracted five genes that are present in RefSeq but that appear to be false (discussed below), to yield 21,306 protein-coding genes (**Table 1**).

**Table 1:**
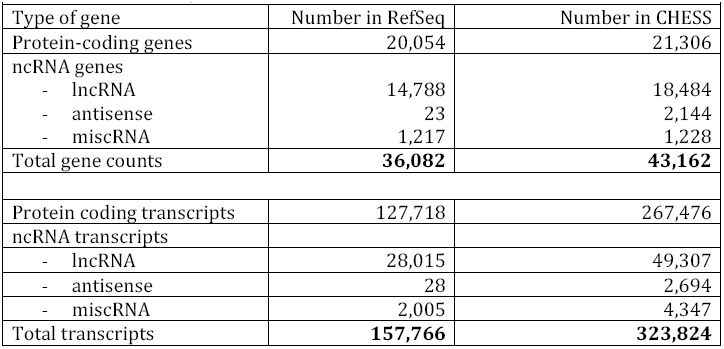
The number of human genes and transcripts in the RefSeq database and in the new CHESS (Comprehensive Human Expressed SequenceS) database built from 9,795 RNA-seq experiments. ncRNA: noncoding RNA; lncRNA: long noncoding RNA gene; miscRNA: miscellaneous RNA.

## Validation using differential expression

As an additional line of evidence that the novel protein-coding genes in CHESS function as genes, we re-analyzed the 9,795 GTEx experiments to test whether any of the novel genes were differentially expressed (DE). If a gene was expressed at significantly different levels–i.e., the transcription level of the gene differed between two conditions–then this finding would support (although not prove) the hypothesis that the gene is genuine.

We conducted two types of tests. First we selected all tissues for which the GTEx data include both male and female samples (21 tissues), and computed which genes were differentially expressed between males and females (see Methods). A total of 207 novel CHESS protein-coding genes were differentially expressed between the sexes (**Figure 2** and Supplementary File S4), and consistent with previously reported results ^28^, breast tissue showed far more DE genes than any other tissue.

**Figure 2:**
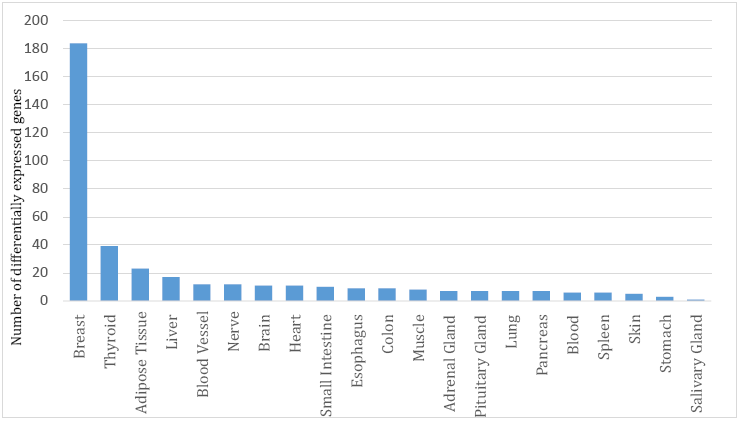
Protein-coding genes that were differentially expressed between males and females, for each of the GTEx tissues that had both male and female samples. All tissues except kidney had at least 10 samples for each sex; kidney had 9 female and 29 male.

Second, we evaluated all genes to determine how many were up-regulated in at least one tissue (see Methods), and found that 998 (84%) of the novel protein-coding genes were up-regulated (**Figure 3** and Suppl. File S5). By comparison, 89% of the RefSeq proteins were up-regulated in at least one tissue. Testis contained the largest number (554) of novel up-regulated genes, with 132 of these genes overlapping retroposons (shown in Suppl. Table S5). While these overlaps might suggest that they are pseudogenes, many retroposed genes have been previously reported as functional, particularly those that exhibit testis-biased expression ^29^.

**Figure 3:**
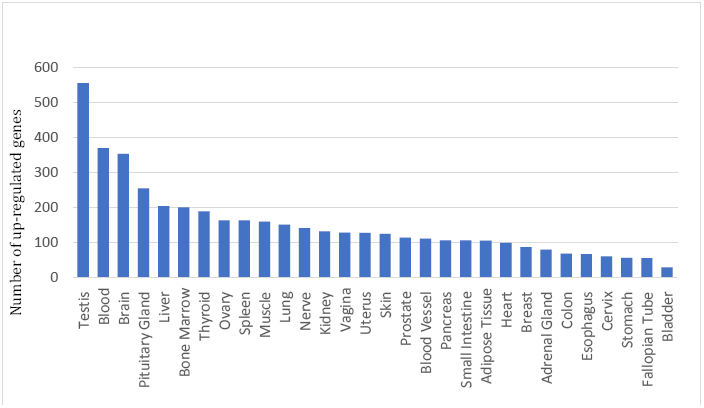
Novel protein-coding genes in CHESS that were up-regulated in the 31 tissues.

## Validation using mass spectrometry

One further possible line of evidence that a locus encodes a protein is direct evidence that the sequence is translated. Publications of two draft human proteomes have recently provided protein evidence for the majority of previously-annotated protein-coding genes, in addition to some previously unknown proteins ^30 31^. These studies and others ^32,33^ suggest that current reference annotation has not yet fully captured the protein-coding potential of the genome. To validate the coding potential of novel loci identified in this study, we searched the unmatched spectra from 30 human tissue/cell types (see Methods) against the novel predicted ORFs described in this study. Peptides identified in this search that were either identical to annotated proteins or mapped with a single mismatch were discarded. We manually examined the MS/MS spectra and discarded those with poor quality. We then created synthetic peptides corresponding to those that supported novel ORFs, and compared the MS/MS spectra from synthetic peptides to experimental spectra.

Based on this analysis pipeline, we identified peptides that confirmed four of the novel protein coding genes in the CHESS set. One example is CHS.57705, a transcript that encodes a 191 amino acid protein that has no similarity to known proteins but is conserved in other primates (**Figure 4a**). This protein contains two transmembrane domains as predicted by SMART ^34^. Another transcript, CHS.24083 encodes a protein of 161 amino acids without any predicted domains or similarity to known proteins (**Figure 4b**), although it too is conserved in primates. Suppl. Table S6 shows all four novel ORFs identified with peptide evidence from proteomics data analysis. Suppl. Figure S12 shows the two additional cases where the mass spectra from synthetic peptides validated the experimental spectra as well as two cases (neither of which passed all the filters required to be a CHESS gene) that were not validated. We note that the abundance of these novel transcripts was very low and the ORFs are relatively short, both of which may explain the small number of identified peptides.

**Figure 4:**
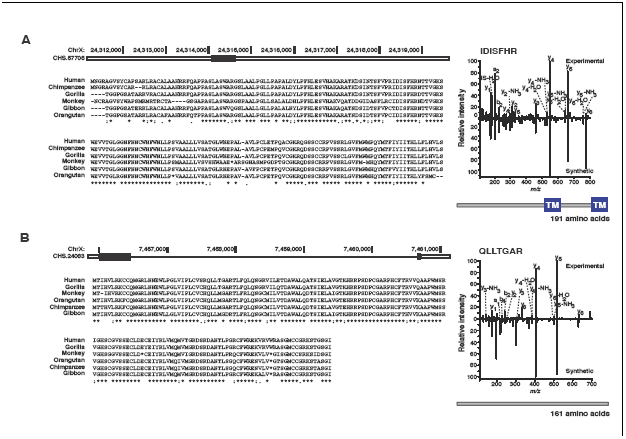
Multiple sequence alignments of novel CHESS protein coding genes CHS.57705 (A) and CHS.24083 (B), each compared to 5 other primates, with annotated MS/MS spectra validating the identified peptides as shown on the right.

## Previously annotated proteins not observed in GTEx

We analyzed the entire set of protein-coding genes in RefSeq to determine how many of them lacked support from any of the 9,795 GTEx samples. We considered a gene to be supported if any GTEx transcript matched any of the gene’s exons; we did not require support for the precise exon-intron structure. Out of all 20,054 RefSeq genes, just ten were not expressed in any of our samples (**Table 2**). We examined each of these ten genes further, and determined that five of them are likely to be errors in RefSeq, as we explain below. We deleted these five genes and their (five) transcripts from the CHESS gene set.

**Table 2:** Protein-coding genes from RefSeq that were not expressed in any of the 9,795 RNA-seq samples from GTeX.

|  |  |  |  |
| --- | --- | --- | --- |
| NCBI Gene ID | Gene name | Location | Product |
| 101927562 | LOC101927562 | chr11<br>1554607–1556457 | uncharacterized |
| 101929097 | LOC101929097 | chr19<br>2511219–2513571 | uncharacterized |
| 107987231 | LOC107987231 | Chr16<br>29973622–29974648 | uncharacterized |
| 101928589 | LOC101928589 | chrX<br>110175773–110177788 | uncharacterized |
| 728072 | CT47A5 | chrX<br>120963026–120966446 | cancer/testis antigen family 47 member A5 |
| 728049 | CT47A8 | chrX<br>120948422–120951842 | cancer/testis antigen family 47 member A8 |
| 728042 | CT47A9 | chrX<br>120943561–120946981 | cancer/testis antigen family 47 member A9 |
| 245927 | DEFB113 | chr6<br>49968677–49969625 | defensin beta 113 |
| 51206 | GP6 | Chr19<br>55013705–55038264 | glycoprotein VI platelet |
| 102723822 | LOC102723822<br>(GTPBP4/NGB) | Unplaced<br>KI270752.1<br>8198–27137 | nucleolar GTP-binding protein 1-like |

The first four genes in Table 3–101927562, 101929097, 107987231, and 101928589–were predicted by computational pipelines at least ten years ago. All loci are entirely contained in the 5’ UTRs of other well-characterized protein-coding genes. GenBank records indicate that the original computational predictions were based on EST evidence and on the presence of open reading frames, but no other evidence supports them. Their position in UTR regions explains the transcript (EST) evidence, but there is no reason to believe these are distinct protein coding genes, and we did not include them in CHESS.

**Table 3:** Genes and transcripts in that are novel in CHESS as compared to RefSeq and Gencode. The columns labelled ‘Annotated in FANTOM’ show the subset of novel CHESS genes and transcripts that are also found in the FANTOM gene catalog.

| **Table 3.** Genes and transcripts in that are novel in CHES as compared to RefSeq and Gencode. The columns labelled ‘Annotated in FANTOM’ show the subset of novel CHES genes and transcripts that are also found in the FANTOM gene catalog. |  |  |  |  |  |  |
| --- | --- | --- | --- | --- | --- | --- |
| Gene biotype | Genes |  |  | Transcripts |  |  |
|  | RefSeq and Gencode | Novel in CHES | Annotated in FANTOM | RefSeq and Gencode | Novel in CHES | Annotated in FANTOM |
| Protein coding | 20,128 | 1,178 | 98 | 169,965 | 97,511 | 23,189 |
| LncRNA | 16,212 | 2,272 | 1,352 | 34,216 | 15,090 | 5,770 |
| Antisense | 598 | 1,546 | 494 | 643 | 2,051 | 606 |
| MiscRNA | 1,227 | 1 | 1 | 2,284 | 2,063 | 476 |

The next three genes in Table 3, CT47A5, CT47A8, and CT47A9, are genes that are normally expressed in germ cells, and reactivated and expressed in some tumors ^35^. Thus it was not surprising that these genes were not expressed in the GTEx samples, which did not include either of these tissue types. Genes DEFB113 and GP6 both appear to be genuine. Both have multiple hits to other proteins, have known functions, and have strong experimental evidence supporting them. It is not clear why they were not present in the GTEx experiments, but it is possible they have highly tissue-specific expression.

Gene 102723822, the final entry in Table 3, is by far the most intriguing of the missing RefSeq proteins. This is a 14-exon gene with a well-characterized product (protein accession XP_006725006), with numerous orthologous proteins in other species. The protein resides on an unplaced scaffold (KI270752) in the current human reference genome, GRCh38.

What is surprising about this protein is that its best alignments are, in order of their similarity, Chinese hamster (*Cricetulus griseus*), golden hamster (*Mesocricetus auratus*), deer mouse (*Peromyscus maniculatus*), Gairdner’s shrewmouse (*Mus pahari*), and other rodents. It is 98% identical to the closest rodent protein, but only 95% identical to the most similar human protein, GTPBP4/NGB on Chr 10 (HGNC:21535). It would be extraordinary for a human protein to have multiple hits to rodents that are all closer than any match to primates. Thus from evolutionary evidence, this protein is clearly a rodent protein, not a human one.

The KI270752 scaffold is 27,745 bp long, and gene 102723822 spans almost all of it, from position 8198 to 27,137. Upon investigation, we discovered that this scaffold is 100% identical to, and clearly derived from, a cosmid deposited in GenBank in 1998, cosmid 1F1 (accession AF065393). We attempted to align the scaffold to an alternate human assembly, CHM1_1.1 (GCA_000306695.2), which was built from whole-genome sequencing of a haploid cell line derived from a human hydatidiform mole, and the scaffold does not match any sequence in that assembly.

Given that K1270752 does not align to any other part of the current human reference genome, that it does not align to an alternative human assembly, and that it contains what appears to be a rodent protein, we concluded that this unplaced scaffold represents contamination in the current human assembly. Thus neither the scaffold nor the gene are human.

We also looked at protein-coding genes that were present in Gencode but not RefSeq. In Gencode release 25, we found 76 genes that were not in RefSeq of which 34 were not expressed in the GTEx experiments (Supplementary File S6). However, in Gencode release 27, all but two of these 34 protein-coding genes were either deleted (27) or changed to noncoding (5), leaving just two genes (AP000351.1 and USP17L23) that were unique to Gencode but not expressed in the GTEx data. Both of these genes are included in the CHESS catalog.

## Non-coding genes

StringTie assembled a total of 30,467,424 transcripts, of which a majority (19,014,285, 62%) had only a single exon (Supplementary Table S1). 1,563,544 transcripts matched RefSeq or Gencode entries, including 209,261 perfect matches and 1,354,283 partial matches. We retained all RefSeq and Gencode transcripts as well as other transcripts for which we found protein-coding evidence, as described above. We then applied a series of filters to remove “noisy” transcripts from the remaining ones, as follows:

1. We required each transcript to be assembled in at least 10 samples, with an average TPM≥1, or alternatively to have expression level as high as the outliers for known transcripts, defined as TPM>13.87 (see Supplementary Materials).
2. We filtered out all single-exon noncoding transcripts.
3. We removed all transcripts that overlapped known LINE elements, LTR repeat elements, or ribosomal RNA genes.
4. To avoid including pre-mRNA transcripts, we removed all transcripts that had retained introns, based on RefSeq and Gencode intron annotations.
5. To eliminate pseudogenes, we filtered out any novel transcript that had at least 98% identity to a known transcript over 90% of its length.
6. We removed all transcripts that overlapped exons of annotated transcripts on the opposite strand, as well as transcripts that overlapped multiple known genes.
7. To reduce transcript assembly false positives, we retained only the ten most abundant novel transcripts at any given locus.
8. We discarded transcripts in loci corresponding to known processed pseudogenes or that overlapped immunoglobulin or T-cell receptor segments.

After applying all the filters above, and including the novel protein coding transcripts described above, we were left with 116,737 transcripts that did not match any RefSeq or Gencode transcripts. Of these, 97,511 represent isoforms (splice variants) of protein-coding genes, increasing the total number of protein-coding transcripts from 127,718 (in RefSeq) to 267,476, or 12.5 isoforms per protein-coding gene (**Tables 1** and **3**). 23,189 of the novel transcripts are also present in the FANTOM database, which used Cap Analysis of Gene Expression (CAGE), to create a large atlas of human genes with high-confidence 5’ ends ^36^.

The number of novel lncRNA gene loci remaining after these filtering steps was 3,819, of which 1,546 were antisense transcripts ^37^, which are contained within introns of other genes. Roughly half of the novel non-coding RNA genes (1,945) were previously also found by the FANTOM consortium ^36^. LncRNA genes have an average of ∼2.6 isoforms in our catalog, although this number could increase if additional evidence emerges in the future.

**Table 1** shows the number of the genes and transcripts, respectively, in protein-coding or long non-coding genes on the GRCh38 human reference sequence. Supplementary Table S3 shows many of the other types of non-coding genes (in addition to lncRNAs) that are annotated in RefSeq. **Table 3** shows the number of genes and transcripts novel to CHESS; i.e., missing in both RefSeq and Gencode.

## Transcriptional noise

Perhaps the most striking result of this study is the vast number of transcripts that appear to have no function at all. Across all data sets and all tissue types, we observed over 30 million distinct transcripts in approximately 700,000 distinct genomic locations, of which only about 40,000 (5%) appear to represent functional gene loci. As others have argued^22^, the mere fact that a sequence is transcribed is insufficient evidence to conclude that it is a gene, despite the fact that early genomics studies made precisely that assumption. It appears instead that 95% of the transcribed locations in the human genome are merely transcriptional noise, explained by the nonspecific binding of RNA polymerase to random or very weak binding sites in the genome. This observation is consistent with efforts to identify sequence motifs that signal the initiation of transcription, which have largely failed because no highly conserved sequences exist.

Similarly, the vast majority of the transcript variants themselves also appear nonfunctional. Although this study greatly increases the number of isoforms of known genes, the 323,824 transcripts reported here represent just 1.1% of the 30,467,424 distinct transcripts observed across all 9,795 data sets. This suggests that the splicing machinery too, like RNA polymerase, is highly nonspecific in its actions, in agreement with previous studies that found that the vast majority of observed splice variants correspond to errors ^38^. The splice sites themselves are much better conserved than any transcription initiation site, but the cellular machinery for cutting and pasting the exons together appears to be inefficient, producing many variations that are simply non-functional, with low abundance isoforms being especially likely to be the result of errors ^39^. It is possible that our criteria for excluding a transcript were too strict, but even so it seems unlikely that a large proportion of the transcripts we rejected are essential for the cell.

Note that functional transcripts occur at much higher abundances than non-functional ones, as shown in Figures S2-S5. If we add up the expression levels of all the functional transcripts and compare that to the total expression of non-functional transcripts, we find that 68% of the transcriptional activity is devoted to producing functional transcripts, while 32% is apparently spent (and presumably wasted) on nonfunctional ones. Thus although the sheer amount of variation is very large, about two-thirds of the RNA molecules in the cell are functional.

## Discussion

The new human gene catalog described here, CHESS, contains a comprehensive set of genes based on nearly 10,000 RNA sequencing experiments. As such, it provides a reference with substantially greater experimental support than previous human gene catalogs. Although it represents only a modest increase in the number of protein-coding genes (1178, or 5.5% out of 21,306 total), it more than doubles the number of splice variants and other isoforms of these genes, to 267,476. This more-comprehensive catalog of genes and splice variants should provide a better foundation for RNA-seq experiments, exome sequencing experiments, genome-wide association studies, and many other studies that rely on human gene annotation as the basis for their analysis.

Given the history of changes in our knowledge of human genes and transcripts, it seems highly likely that this new database will change further in the future. In particular, many of the more than 18,000 noncoding RNA genes have less evidential support than the protein-coding component of the genome, and this number may decline over time just as the human gene count declined from 2001 to the present. The CHESS database of genes and transcripts, which is freely available at *http://ccb.jhu.edu/chess*, will be updated over time as new evidence emerges.

The overall picture that emerges from this analysis is that the cell is a relatively inefficient machine, transcribing more DNA into RNA than it needs. Ever since the discovery of introns ^40,41^, we have known that genomes contain large regions that appear to have no function.

Based on the results described here, it appears that nearly 99% of the transcriptional variety produced in human cells has no apparent function, although most of these variants appear at such low levels that they cumulatively account for only 32% of transcriptional volume.

## Methods

The initial GTEx data release contained 1641 RNA-seq samples ^20,42^, and a subsequent publication described a much larger set of 8555 samples collected from 40 body sites ^43^. Our data represents a later GTEx data release with 9,795 samples across 31 tissue types and 54 body sites, summarized in Supplementary Table S2. The GTEx data used for the analyses described in this study were obtained from dbGaP accession number phs000424.v6.p1 in May of 2016.

*Alignment and assembly*. In total, the 9,795 RNA-seq samples contain 899,960,113,026 reads (449,980,056,513 pairs), an average of 91.9 million reads (46M pairs) per sample. The RNA-seq assembly process, illustrated in **Figure 5**, requires multiple steps of alignment, assembly, and quantification ^44^ for each of the samples. We aligned each sample to release GRCh38.p8 of the human genome using HISAT2 ^45^ with default parameters, providing it with the RefSeq annotation. We then assembled the alignments using StringTie ^46^ again providing the RefSeq annotation. Both HISAT2 and StringTie use annotation as a guide when provided, but both programs find novel splice sites (HISAT) and novel transcripts (StringTie) whenever necessitated by the data. The RefSeq annotation provided here contained 20,054 protein-coding genes, 15,779 long noncoding RNA (lncRNA) genes, 16,131 pseudogenes, and 629 tRNA genes, as well as a few other specialized categories of annotation (Supplementary Table S3).

**Figure 5:**
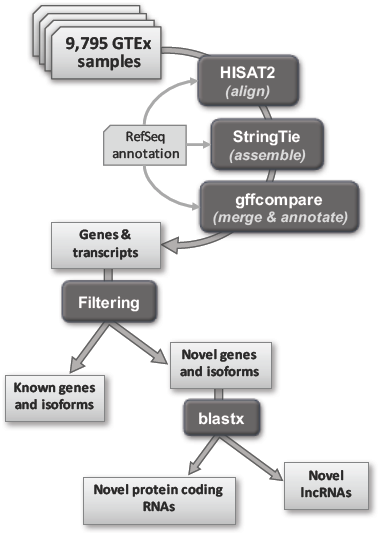
Summary of computational pipeline used to align and assemble all 9,795 RNA-seq samples.

*Timings.* Alignment of reads with HISAT2 was the most computationally intensive step in the process. On average, 88.3% of reads aligned successfully across all 9,795 samples. The alignment steps included uncompressing the original SRA files, aligning them to the genome with HISAT2 (using 8 CPUs in parallel), and compressing the output to produce BAM files. These steps took an average of 43 minutes per file (sample), using a 32-core server with a 2.13 GHz Intel Xeon E7 for benchmarking. Following alignment, the aligned reads were sorted and converted to the compact CRAM format. This process took an average of 17 minute per sample using 8 threads. Assembly and quantification with StringTie took an average of 24 minutes per sample using 4 threads. Thus the total average time to process on sample, with much of the time limited by I/O speed for the very large files involved, was 84 minutes. Processing all 9,795 samples required about 13,700 hours (571 days); by dividing the computation across many processors this was reduced to about 30 days total elapsed time. Note that attempts to parallize this process further would require distributing the files across many independent storage units; otherwise contention for file access would make parallel processing ineffective.

After the initial assembly steps, many transcripts were fragmented (i.e., not full-length) due to low coverage in particular samples. To correct this problem, we compared all transcripts and transcript fragments across all samples, and merged any transcripts that were contained or overlapped with others. For this merging step, we first used the program gffcompare (ccb.jhu.edu/software/stringtie/gff.shtml) to merge all GTF (Gene Transfer Format) files from the original samples on a tissue-by-tissue basis. Following this step, which produced a single GTF file for each of the 31 tissues, we merged the 31 files together to produce a single, consistent set of transcripts that accounted for all samples.

We computed expression levels using both TPM (transcripts per million reads) and FPKM (fragments per kilobase of exon per million reads). The Supplementary Materials file contains detailed statistics on expression levels for all genes in RefSeq, including distributions used to calculate outliers for protein coding and lncRNA genes (Figures S1– S5. For consistency, we used TPM values as thresholds for all filtering steps.

We identified the longest open reading frames (ORFs) in transcripts with gffread (github.com/gpertea/gffread). We ran BLAST searches of all ORFs against the Swiss-Prot section of UniProt (release 2017_09) and the **nr** database, a comprehensive, non-redundant protein database downloaded from NCBI in February 2017. The mammalian protein database used in some searches was a subset of nr. When considering lncRNA matches to proteins from this database, we considered a protein to have unknown function if its name included any of the following keywords: hypothetical, unnamed protein product, uncharacterized protein, unknown, pseudogene, LOC, PRO, orf, or open reading frame. We also excluded proteins whose only annotation was a name with the prefix hCG, which are computational predictions based only on *de novo* gene finding programs and/or EST evidence reported in one of the original human genome papers ^7^. The multiple alignment shown in Figure 1 was produced with SeaView v4.6.2 ^47^.

*Comparisons to known annotation databases*. We used gffcompare to compare assembled transcripts to the RefSeq and GENCODE databases. We downloaded the FANTOM transcripts defined as ‘robust’ from fantom.gsc.riken.jp/5/suppl/Hon_et_al_2016/data/assembly/lv3_robust/ and used the UCSC’s liftOver program with the default parameters to remap them from GRCh37 to GRCh38. We checked all ∼30 million StringTie-assembled transcripts against these remapped FANTOM transcripts using the trmap program (github.com/gpertea/trmap), a specialized version of gffcompare, optimized for streaming a large set of transcripts against another set, for the purpose of reporting and classifying overlaps between them.

Because the FANTOM transcripts had experimental data to support their 5’ ends, we adjusted the ends of the CHESS transcripts when they otherwise matched the full length of a FANTOM transcript.

*Differential expression between sexes*. We used Salmon ^48^ to generate quantification estimates of the complete set of CHESS transcripts assembled from the GTEx data. The advantages to using Salmon over other transcript quantification programs include speed, compatibility with downstream analysis tools, and the ability to retain multi-mapped reads. Salmon relies on the use of an index built from transcript sequences to quasi-map RNA-seq reads in the quantification step. To obtain these sequences, we used gffread (http://ccb.jhu.edu/software/stringtie/gff.shtml) to extract them from the CHESS GFF file. The index built from the resulting multi-fasta file and the raw sequencing reads were then used to generate CHESS transcript abundance estimates for each GTEx sample.

We used the tximport package ^49^ to import the Salmon output and generate separate gene-level count matrices for each tissue that contained both male and female samples. To account for the widely varying number of samples per tissue, we chose a random subset of samples from tissues with large numbers of samples.

We then used the resulting count matrices as input to DESeq2 ^50^ to conduct differential expression analysis within each of the 31 tissues independently, comparing male to female samples. We used the False Discovery Rate (FDR) computed from DESeq2’s implementation of the Benjamini-Hochberg adjustment. The set of differentially expressed genes with FDR < 0.05 was then filtered to identify the protein-coding genes that were novel in CHESS.

From the results of the 31 DESeq2 experiments, we created a single list of genes differentially expressed in at least one tissue. For breast tissue we counted all genes with a FDR < 0.05. For each additional tissue, we only included genes with an FDR < 0.002 to correct for multiple comparisons across tissues. An FDR threshold of 0.002 for each gene in each tissue results in an FDR of ∼0.05 for each gene across all tissues. The major differences between male and female breast tissue lead us to expect a large number of differentially expressed genes.

*Tissue-specific differential expression.* We started by randomly selecting 20 samples from each of the 31 tissues. In cases where the given tissue had fewer than 20 samples, we selected all samples. Using tximport, we then created one gene-level count matrix for these 591 samples. With this count matrix, we ran DESeq2 to test for differential expression between tissues while controlling for the effect of sex. Using the ‘contrast’ argument of the results function in DESeq2, we made 31 different comparisons to find genes up-regulated in each tissue. Each comparison contrasted the gene expression in the tissue of interest to the average expression across all other tissues. For each tissue, we considered up-regulated genes with an FDR < 0.05 significant and then filtered this list to report only novel, protein coding genes. To create a list of all novel genes up-regulated in at least one tissue, we reduced the FDR threshold to 0.0015 to correct for the 31 comparisons. An FDR threshold of 0.0015 for each gene in each tissue results in a (conservative) FDR of ∼0.05 for each gene across all tissues.

*Mass spectrometry*. Unmatched MS/MS spectra from a previous study ^30^ were searched against translated products of predicted CHESS ORFs using the SEQUEST search engine on Proteome Discoverer 2.1 software platform (Thermo Fisher Scientific).

Carbamidomethylation of cysteine and oxidation of methionine were specified as fixed and variable modifications. Mass tolerance limits were set to 10 ppm and 0.02 Da for precursor and fragment ions, respectively. A target-decoy database approach was employed to filter the identified peptides at a 1% false discovery rate. Peptide sequences that corresponded to novel genes were synthesized (JPT Peptide Technologies, Berlin, Germany), analyzed on an Orbitrap Fusion Lumos Tribrid mass spectrometer (Thermo Fisher) and compared against the experimental spectra. Putative translational products of novel ORFs were aligned using BLAST against the NCBI ‘nr’ protein database and domain prediction was carried out using SMART ^34^. Multiple sequence alignment of protein sequences was performed using Clustal Omega ^51^.

## Acknowledgements

Thanks to Tajer Mun for help with BLAST protein searches. This work was supported in part by NSF grant DBI-1458178 to M.P., NIH grants R01-HG006677 and R01-GM083873 to S.L.S., and NIH grant U24-CA210985 to A.P. The GTEx Project was supported by the Common Fund of the Office of the Director of the National Institutes of Health, and by NCI, NHGRI, NHLBI, NIDA, NIMH, and NINDS.

## References

1. Vogel, F. A Preliminary Estimate of the Number of Human Genes. Nature 201, 847 (1964).

2. Schuler, G.D. et al. A gene map of the human genome. Science 274, 540–6 (1996).

3. Antequera, F. & Bird, A. Predicting the total number of human genes. Nat Genet 8, 114 (1994).

4. Fields, C., Adams, M.D., White, O. & Venter, J.C. How many genes in the human genome? Nat Genet 7, 345–6 (1994).

5. Liang, F. et al. Correction: Gene Index analysis of the human genome estimates approximately 120,000 genes. Nature Genetics 26, 501 (2000).

6. The International Human Genome Sequencing Consortium. Initial sequencing and analysis of the human genome. Nature 409, 860–921 (2001).

7. Venter, J.C. et al. The sequence of the human genome. Science 291, 1304–51 (2001).

8. International Human Genome Sequencing Consortium. Finishing the euchromatic sequence of the human genome. Nature 431, 931–45 (2004).

9. Clamp, M. et al. Distinguishing protein-coding and noncoding genes in the human genome. Proc Natl Acad Sci U S A 104, 19428–33 (2007).

10. Ezkurdia, I. et al. Multiple evidence strands suggest that there may be as few as 19,000 human protein-coding genes. Hum Mol Genet 23, 5866–78 (2014).

11. Pertea, M. & Salzberg, S.L. Between a chicken and a grape: estimating the number of human genes. Genome Biol 11, 206 (2010).

12. O’Leary, N.A. et al. Reference sequence (RefSeq) database at NCBI: current status, taxonomic expansion, and functional annotation. Nucleic Acids Res 44, D733–45 (2016).

13. Harrow, J. et al. GENCODE: the reference human genome annotation for The ENCODE Project. Genome Res 22, 1760–74 (2012).

14. Farrell, C.M. et al. Current status and new features of the Consensus Coding Sequence database. Nucleic Acids Res 42, D865–72 (2014).

15. Need, A.C. et al. Clinical application of exome sequencing in undiagnosed genetic conditions. J Med Genet 49, 353–61 (2012).

16. Zhu, X. et al. Whole-exome sequencing in undiagnosed genetic diseases: interpreting 119 trios. Genet Med 17, 774–81 (2015).

17. Guttman, M. et al. Chromatin signature reveals over a thousand highly conserved large non-coding RNAs in mammals. Nature 458, 223–7 (2009).

18. Cabili, M.N. et al. Integrative annotation of human large intergenic noncoding RNAs reveals global properties and specific subclasses. Genes Dev 25, 1915–27 (2011).

19. Kung, J.T., Colognori, D. & Lee, J.T. Long noncoding RNAs: past, present, and future. Genetics 193, 651–69 (2013).

20. The GTEx Consortium. Human genomics. The Genotype-Tissue Expression (GTEx) pilot analysis: multitissue gene regulation in humans. Science 348, 648–60 (2015).

21. Adams, M.D., Kerlavage, A.R., Fields, C. & Venter, J.C. 3,400 new expressed sequence tags identify diversity of transcripts in human brain. Nat Genet 4, 256–67 (1993).

22. Palazzo, A.F. & Lee, E.S. Non-coding RNA: what is functional and what is junk? Front Genet 6, 2 (2015).

23. Raj, A., Peskin, C.S., Tranchina, D., Vargas, D.Y. & Tyagi, S. Stochastic mRNA synthesis in mammalian cells. PLoS Biol 4, e309 (2006).

24. Trapnell, C. et al. Transcript assembly and quantification by RNA-Seq reveals unannotated transcripts and isoform switching during cell differentiation. Nat Biotechnol 28, 511–5 (2010).

25. Trapnell, C. et al. Differential gene and transcript expression analysis of RNA-seq experiments with TopHat and Cufflinks. Nat Protoc 7, 562–78 (2012).

26. Altschul, S.F. et al. Gapped BLAST and PSI-BLAST: a new generation of protein database search programs. Nucleic Acids Res 25, 3389–402 (1997).

27. The UniProt, C. UniProt: the universal protein knowledgebase. Nucleic Acids Res 45, D158–D169 (2017).

28. Mele, M. et al. Human genomics. The human transcriptome across tissues and individuals. Science 348, 660–5 (2015).

29. Emerson, J.J., Kaessmann, H., Betran, E. & Long, M. Extensive gene traffic on the mammalian X chromosome. Science 303, 537–40 (2004).

30. Kim, M.S. et al. A draft map of the human proteome. Nature 509, 575–81 (2014).

31. Wilhelm, M. et al. Mass-spectrometry-based draft of the human proteome. Nature 509, 582–7 (2014).

32. Na, C.H. et al. Discovery of noncanonical translation initiation sites through mass spectrometric analysis of protein N termini. Genome Res 28, 25–36 (2018).

33. Samandi, S. et al. Deep transcriptome annotation enables the discovery and functional characterization of cryptic small proteins. Elife 6(2017).

34. Letunic, I. & Bork, P. 20 years of the SMART protein domain annotation resource. Nucleic Acids Res 46, D493–D496 (2018).

35. Chen, Y.T. et al. Identification of a new cancer/testis gene family, CT47, among expressed multicopy genes on the human X chromosome. Genes Chromosomes Cancer 45, 392–400 (2006).

36. Hon, C.C. et al. An atlas of human long non-coding RNAs with accurate 5’ ends. Nature 543, 199–204 (2017).

37. Mercer, T.R., Dinger, M.E., Sunkin, S.M., Mehler, M.F. & Mattick, J.S. Specific expression of long noncoding RNAs in the mouse brain. Proc Natl Acad Sci U S A 105, 716–21 (2008).

38. Saudemont, B. et al. The fitness cost of mis-splicing is the main determinant of alternative splicing patterns. Genome Biol 18, 208 (2017).

39. Pickrell, J.K., Pai, A.A., Gilad, Y. & Pritchard, J.K. Noisy splicing drives mRNA isoform diversity in human cells. PLoS Genet 6, e1001236 (2010).

40. Chow, L.T., Gelinas, R.E., Broker, T.R. & Roberts, R.J. An amazing sequence arrangement at the 5’ ends of adenovirus 2 messenger RNA. Cell 12, 1–8 (1977).

41. Berget, S.M., Moore, C. & Sharp, P.A. Spliced segments at the 5’ terminus of adenovirus 2 late mRNA. Proc Natl Acad Sci U S A 74, 3171–5 (1977).

42. Carithers, L.J. et al. A Novel Approach to High-Quality Postmortem Tissue Procurement: The GTEx Project. Biopreserv Biobank 13, 311–9 (2015).

43. Wheeler, H.E. et al. Survey of the Heritability and Sparse Architecture of Gene Expression Traits across Human Tissues. PLoS Genet 12, e1006423 (2016).

44. Pertea, M., Kim, D., Pertea, G.M., Leek, J.T. & Salzberg, S.L. Transcript-level expression analysis of RNA-seq experiments with HISAT, StringTie and Ballgown. Nat Protoc 11, 1650–67 (2016).

45. Kim, D., Langmead, B. & Salzberg, S.L. HISAT: a fast spliced aligner with low memory requirements. Nat Methods 12, 357–60 (2015).

46. Pertea, M. et al. StringTie enables improved reconstruction of a transcriptome from RNA-seq reads. Nat Biotechnol 33, 290–5 (2015).

47. Gouy, M., Guindon, S. & Gascuel, O. SeaView version 4: A multiplatform graphical user interface for sequence alignment and phylogenetic tree building. Mol Biol Evol 27, 221–4 (2010).

48. Patro, R., Duggal, G., Love, M.I., Irizarry, R.A. & Kingsford, C. Salmon provides fast and bias-aware quantification of transcript expression. Nat Methods 14, 417–419 (2017).

49. Soneson, C., Love, M.I. & Robinson, M.D. Differential analyses for RNA-seq: transcript-level estimates improve gene-level inferences. F1000Res 4, 1521 (2015).

50. Love, M.I., Huber, W. & Anders, S. Moderated estimation of fold change and dispersion for RNA-seq data with DESeq2. Genome Biol 15, 550 (2014).

51. Sievers, F. et al. Fast, scalable generation of high-quality protein multiple sequence alignments using Clustal Omega. Mol Syst Biol 7, 539 (2011).

